# Silencing of oncogenic KRAS by mutant-selective small interfering RNA

**DOI:** 10.1101/2020.10.08.331835

**Authors:** Bjoern Papke, Salma H. Azam, Anne Y. Feng, Amanda E. D. Van Swearingen, Christina Gutierrez-Ford, Pradeep S. Pallan, Martin Egli, Adrienne D. Cox, Channing J. Der, Chad V. Pecot

## Abstract

Oncogenic mutations in the *KRAS* gene are well-established drivers of cancer. While the recently developed KRAS^G12C^ inhibitors offer a targeted KRAS therapy and have shown success in the clinic, KRAS^G12C^ represents only 11% of all KRAS mutations. Current therapeutic approaches for all other KRAS mutations are both indirect and non-mutant-selective, largely focusing on inhibition of downstream KRAS effectors such as MAP kinases. Inhibition of KRAS downstream signaling results in a system-wide down-modulation of the respective targets, raising concerns about systemic cell toxicity. Here, we describe a custom short interfering RNA (siRNA) oligonucleotide (EFTX-D1) designed to preferentially bind mRNA of the most commonly occurring KRAS missense mutations in codons 12 and 13. We determined that EFTX-D1 preferentially reduced the mutant KRAS sequence versus wild-type at the levels of both transcription and translation, and reversed oncogenic KRAS-induced morphologic and growth transformation. Furthermore, EFTX-D1 significantly impaired the proliferation of several KRAS mutant cancer cell lines in 2-D as well as 3-D assays. Taken together, our data indicate a novel use of RNA interference (RNAi) to target oncogenic KRAS-driven cancers specifically.

## Introduction

The KRAS oncogene is amongst the most frequent and lethal drivers of cancers in the world and has a mutation frequency of approximately 31% in lung, 45% in colorectal, and 98% in pancreatic cancers (Cox et al. 2014; Siegel, Miller, and Jemal 2020). Mutations in one of three distinct hot spot codons (12, 13, or 61) impair KRAS GTPase activity and thereby lock it in a GTP-bound active state (Hobbs, Der, and Rossman 2016). In the active state, KRAS continuously stimulates downstream effector signaling, resulting in increased proliferation and cell survival (Stephen et al. 2014).

While the KRAS oncogene is considered a primary drug target in anti-cancer drug discovery, KRAS poses unique structural and functional challenges which have made this protein particularly difficult to target. For example, attempts to develop GTP-competitive inhibitors of RAS (inspired by the success of ATP-competitive inhibitors for other targets) have failed due to the many-fold higher nanomolar affinity of GTP for KRAS, which is difficult to outcompete (Cox et al. 2014). Structural analysis of the KRAS protein has revealed that there are no deep hydrophobic pockets readily available for binding, making the development of potent small molecule inhibitors challenging (Grant et al. 2011; Cox et al. 2014). Alternatively, attempts to inhibit key downstream effectors of KRAS, predominantly in the RAF−MEK−ERK and the PI3K−AKT−mTOR pathways, have been tempered by the significant crosstalk between pathways, activation of compensatory mechanisms, and functional redundancy (i.e., multiple isoforms and pathways can contribute to tumor progression) (Papke and Der 2017; Waters and Der 2018). Thus, combination approaches to inhibit multiple pathways are needed to effectively shut down KRAS oncogenic activity (Engelman et al. 2008); however, this raises concerns for widespread inhibition of other pathways and resultant unwanted toxicity (Cox et al. 2014).

The first true advancement in direct KRAS inhibition has been made only recently with the development of novel drugs currently in clinical trials, including AMG510 and MRTX849, which covalently bind to KRAS^G12C^ to lock it in its inactive GDP-bound state (Janes et al. 2018; Ostrem et al. 2013; Canon et al. 2019; Hallin et al. 2020). However, this is unlikely to be a silver bullet approach for all KRAS mutations, as G12C presents some vulnerabilities unique to this mutation. For example, the lysine residue (Lys-16) in the binding pocket catalyzes covalent binding, and the G12C protein has one of the fastest turnover rates of all the KRAS mutants, allowing more frequent drug occupancy in the inactive GDP-bound state (McCormick 2020). Therefore, although the recent advance of these G12C inhibitors has aroused unprecedented excitement in the KRAS community, the need for effective inhibition strategies for the other predominant KRAS mutations remains a critical unmet need, given that G12C accounts for only 11% of all KRAS mutant cancers. In addition, it is important to note that even for G12C, the success of these inhibitors is still tempered by limited efficacy in the clinic, due in part to the emergence of resistance mechanisms. For example, some cells that enter quiescence upon KRAS G12C inhibition can stimulate a compensatory increase in production of new KRAS G12C protein that maintains its active (GTP-bound), drug-insensitive state, thereby evading drug inhibition (Xue et al. 2020). Thus, an alternative strategy that inhibits protein production (as opposed to protein function), such as RNA interference (RNAi), may be an efficacious approach that abrogates this type of resistance to targeting KRAS. Of note, efforts to develop PROTACs against KRAS G12C in lung and pancreatic cancer have been shown to fail due to shortcomings such as an inability to successfully promote poly-ubiquitination of endogenous KRAS G12C (Zeng et al. 2020).

Therapeutic RNAi refers to the delivery into cells of synthetic oligonucleotides complementary to specific target RNAs, which then utilize the endogenous RNA-induced silencing complex (RISC) to cleave those messenger RNA (mRNA) sequences or to inhibit their translation (Fire et al. 1998). Previous work from our lab and others has investigated the possibility of therapeutically down-modulating KRAS mRNA expression via siRNA (Yuan et al. 2014; Pecot et al. 2014; Xue et al. 2014) or antisense oligonucleotides (Ross et al. 2017). However, one shortcoming of these approaches is their lack of mutant versus wild-type (WT) specificity. Systemic equal silencing of both mutant and WT KRAS mRNAs may give rise to toxicity due to the loss of KRAS WT in healthy tissues. Although the consequences of toxicity following KRAS loss in healthy adult tissues remain to be evaluated, selective inhibition of tumor-specific mutant KRAS could have significant advantages for an optimal therapeutic window. Here we report the design and development of EFTX-D1, a mutant-selective, WT-sparing siRNA that has a higher affinity for several of the most common oncogenic KRAS mutations compared to WT KRAS.

## Results

### EFTX-D1 is designed to target the most common KRAS codon 12 and codon 13 mutations but spare wild-type KRAS

Codon 12 and codon 13 mutations are the most frequently occurring KRAS mutations in cancer, and G12D, G12V, G12C and G13D account for about 80% of all KRAS mutations (Fig. 1a). Due to the close proximity of these mutational hotspots, we sought to determine whether an anti-sense sequence of customized siRNAs could target several of these mutations yet spare the wild-type sequence. Using the Block-iT RNAi Designer tool (Invitrogen), we designed a library of siRNAs that target an engineered, artificial KRAS mRNA sequence that simultaneously contains the point mutations of G12C, G12D, and G13D (Fig. 1b, c). We termed the siRNAs targeting this sequence EFTX-D siRNAs. Because G12D and G12V missense mutations occur in the same nucleotide of codon 12, we also engineered an artificial KRAS mRNA sense sequence carrying the G12C, G12V, and G13D mutations, which served as a target for EFTX-V siRNAs (Fig. 1b, Supplementary Fig. 1). Previous literature suggests there is a 3-nucleotide mismatch tolerance threshold for 19 base pair siRNA efficacy of mRNA target recognition and knockdown (Naito et al. 2004). EFTX-D and EFTX-V siRNAs were designed to have only 2 mismatches with any of their target KRAS mutation sequences (e.g. EFTX-D siRNAs have 2 mismatches with a KRAS G12C, G12D, or G13D mRNA transcript), while siRNA binding to the WT KRAS mRNA contains 3 mismatches (Fig 1c, Supplementary Fig. 1). We then evaluated whether this imbalance of 2 versus 3 mismatches was sufficient to create preferential inhibition of the target KRAS mutation while sparing the WT sequence.

**Fig 1.**
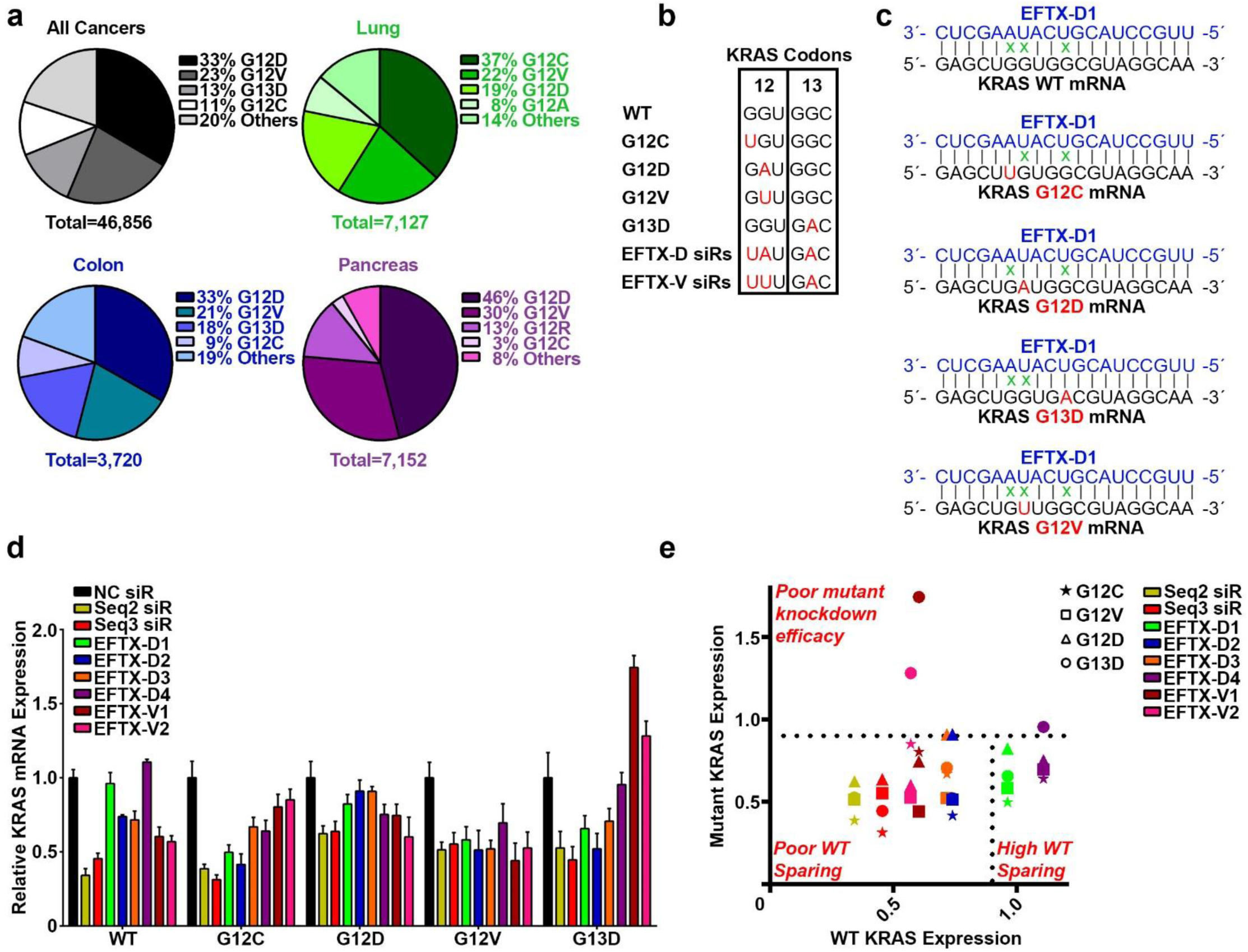
Design and selection of a KRAS-mutant, WT-sparing siRNA. **a** Most common KRAS mutations in all cancers (grey), lung cancer (green), colon cancer (blue) and pancreatic cancer (purple). **b** Sequence alignment of KRAS codons 12 and 13 in the wild-type (WT) form with the 4 most commonly mutated variants (G12C, G12D, G12V, and G13D) as well as EFTX-D siRNAs (siRs) and EFTX-V siRs (with each class of siRs targeting a hypermutated KRAS sequence carrying 3 oncogenic mutations). Mutated nucleotides are in red. **c** Pairing of the EFTX-D1 siR antisense strand with KRAS WT, G12C, G12D, G12V, and G13D mRNA transcripts. Non-Watson-Crick base pairings (i.e., mismatches) are indicated by a green *x*. **d** qPCR of NIH/3T3 cells stably expressing KRAS WT, G12C, G12D, G12V, or G13D and transiently transfected with negative control (NC), Seq2, Seq3, EFTX-D or EFTX-V siRs at a dose of 20 nM. Cells were analyzed 24 hrs post transfection and qPCR was run in technical triplicate. Results were normalized to NC siR transfection. **e** Revisualization of (**d**) by comparing level of WT expression (x-axis) vs expression of each KRAS mutant (y-axis) after transfection of each siR assessed in (**d**). Results in the top half above the horizontal dotted line indicate poor knockdown efficacy of mutant KRAS. Results in the bottom left indicate effective knockdown of mutant KRAS but poor WT sparing (i.e., concomitant knockdown of mutant and WT KRAS expression). Lead siRs appear in the bottom right quadrant and demonstrate potent mutant inhibition and optimal WT sparing. Symbol color indicates siR treatment; symbol shape indicates KRAS mutation assessed. Results were normalized to NC siR transfection.

**Fig. 2.**
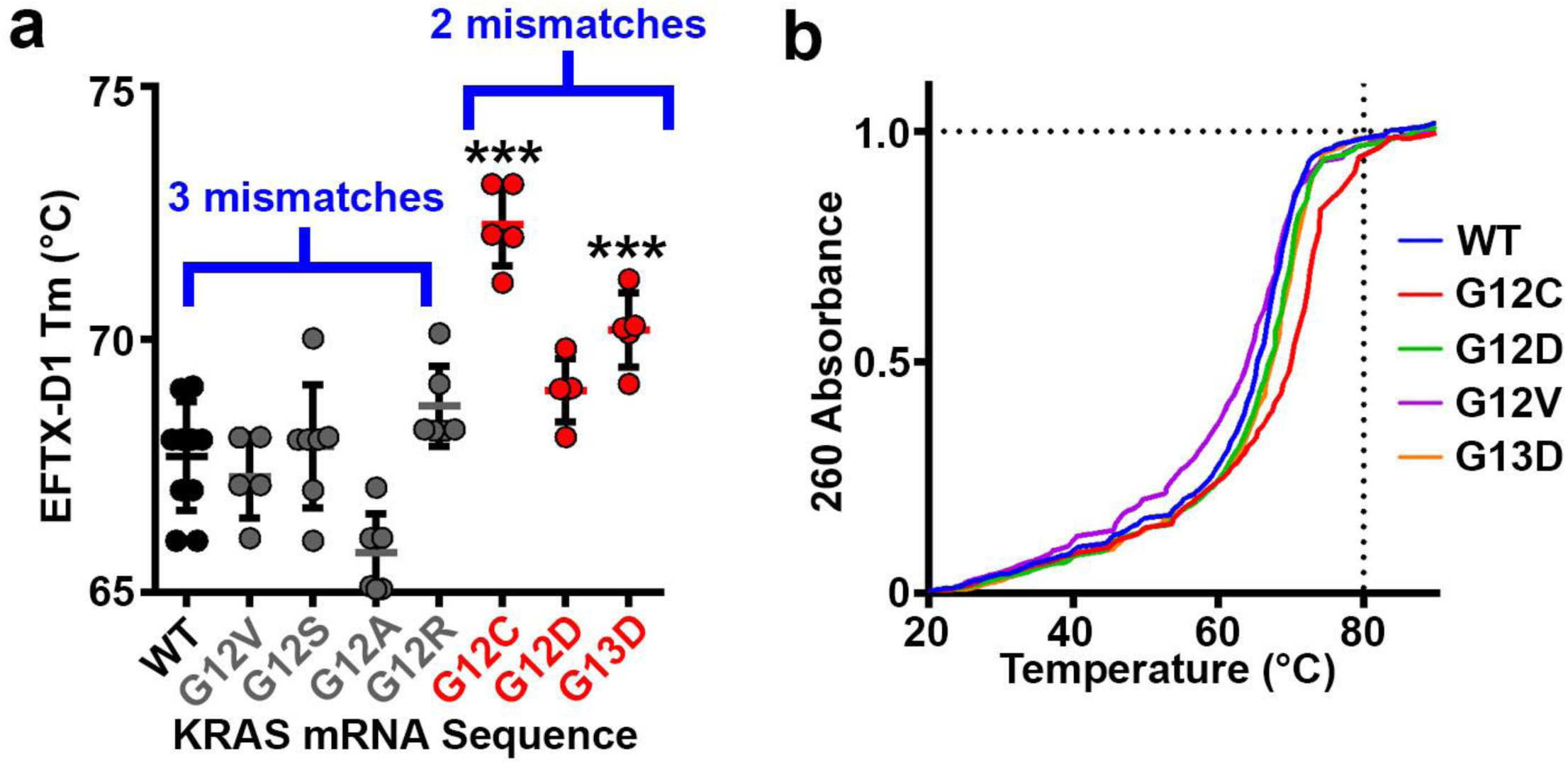
Thermodynamic stability of EFTX-D1 and KRAS mRNA association. **a** Melting temperatures T_m_ of EFTX-D1 siRNA paired with each indicated target KRAS mRNA synthetic sequence. WT and grey mutants carry 3 mismatches with EFTX-D1; red mutants carry 2 mismatches with EFTX-D1. **b** Melting curves of EFTX-D1 with each indicated target KRAS mRNA synthetic sequence. Thirteen technical replicates were run for WT and 6-7 replicates for each mutant. Statistical significance was measured by a one-way ANOVA test; p-values are indicated as ***p<0.001.

Candidate EFTX-D and EFTX-V siRNAs were transiently transfected into NIH/3T3 mouse fibroblasts stably expressing human KRAS WT, G12C, G12D, G12V, or G13D, and real-time qPCR was used to screen for their ability to potently inhibit expression of mutant KRAS while sparing WT (Fig. 1d). Knockdown efficacy was compared to those of our previously published pan-KRAS siRNA sequences Seq2 and Seq3. These bind to a non-mutated, conserved region of KRAS mRNA far downstream of codon 13 (0 mismatches with G12 and G13 mutants, and with WT), rendering them non-specific for mutant versus WT KRAS. We have previously demonstrated that Seq2 and Seq3 potently inhibit KRAS expression *in vitro*, and exert therapeutic inhibition of KRAS-driven tumor progression in lung and colon cancer models (Pecot et al. 2014). Both EFTX-D1 and EFTX-D4 demonstrated high mutant-specificity and potency of knockdown relative to negative control (NC) siRNA. However, EFTX-D1 exhibited a superior ability to consistently inhibit expression of the targeted mutants while maintaining WT sparing (Fig. 1e). Unexpectedly, EFTX-D1 also demonstrated activity against the G12V mutant, which has 3 base pair mismatches (Fig. 1d,e). Six additional EFTX-D siRNA sequences, with all possible starting positions between EFTX-D1 and EFTX-D4, were also tested for their ability to reduce mutant KRAS, but all proved to be inferior to the KRAS silencing potency of EFTX-D1 (Supplementary Fig. 3).

**Fig. 3.**
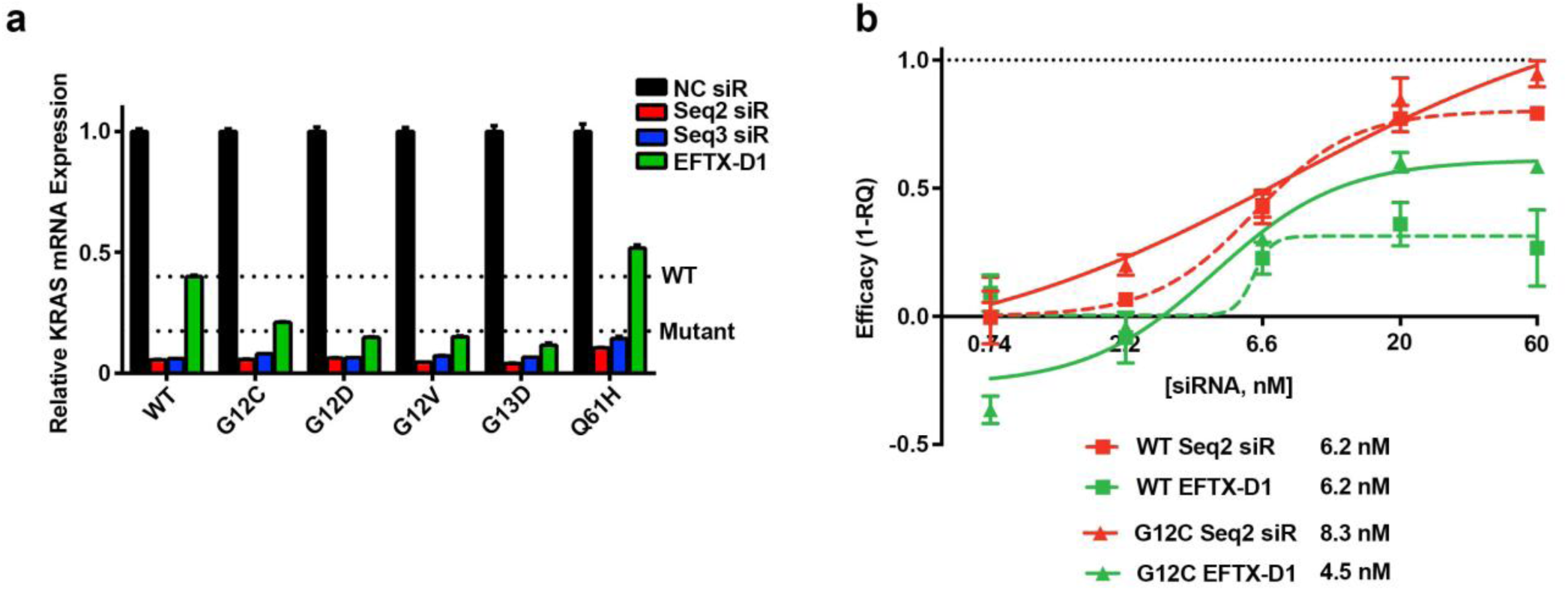
KRAS mutant-specific knockdown by EFTX-D1. **a** qPCR at 24 hours post transfection of NIH/3T3 cells stably expressing KRAS WT, G12C, G12D, G12V, G13D, or Q61H and transiently transfected with 40 nM NC, Seq2, Seq3, or EFTX-D1 siRNA (siR). The “WT” line indicates average expression level of KRAS mRNA with WT codon 12 and codon 13 sequences, in response to EFTX-D1 transfection. The “Mutant” line indicates average expression level of the KRAS mutants carrying codon 12 or codon 13 mutations, in response to EFTX-D1 transfection. qPCR was run in duplicate. **b** qPCR knockdown efficacy dose response curves of NIH/3T3 cells stably expressing KRAS WT or G12C and transiently transfected with Seq2 or EFTX-D1 siR. Results are shown 24 hrs post transfection and are normalized to NC siR transfection. qPCR was run in duplicate. IC_50_ values for siR treatment of each cell line are shown below.

### WT KRAS sparing relative to mutant KRAS targeting is due to reduced thermodynamic stability of the WT KRAS-EFTX-D1 complex

We predicted that the WT sparing feature of EFTX-D1 would be due to differences in the binding energy of EFTX-D1 to its 2-mismatch mutant target transcripts versus the 3-mismatch WT target. To test this possibility, we characterized the melting points of EFTX-D1 complexed with different synthesized KRAS mRNA mimics (Fig. 2a, b). This allowed us to determine the binding strength between WT or mutant KRAS mRNA and EFTX-D1. We observed that the average melting temperature was lower (reflecting reduced thermodynamic stability of binding) for pairing reactions containing KRAS WT (67.7°C) or KRAS mutations with 3 mismatches (G12V – 67.3°C, G12S – 67.9°C, G12A – 65.8°C, and G12R – 68.7°C) compared to the reactions containing KRAS mutations with 2 mismatches on EFTX-D1 (G12C – 72.3°C, G12D – 69.0°C, and G13D – 70.2°C).

### EFTX-D1 selectively reduces oncogenic KRAS mRNA

To further characterize whether EFTX-D1 has a significant difference in downmodulation of mutant, oncogenic KRAS mRNA over WT KRAS mRNA, we performed transient knockdown experiments in our NIH/3T3 cell lines stably expressing human KRAS WT, G12C, G12D, G12V, or the codon 61 mutation Q61H (Fig. 3a). Whereas EFTX-D1 has only a 2 base pair mismatch with G12C and G12D, and a 3 base pair mismatch with WT and G12V, it also has a 3 base pair mismatch with the Q61H mutant, as the nucleotide sequence of codons 12 and 13 in this mutant is the same as WT. We therefore anticipated that EFTX-D1 would downmodulate Q61H mRNA to the same extent as WT mRNA. Comparing the KRAS knockdown efficacy of EFTX-D1 to that of the pan KRAS (i.e., both WT and mutant-targeting) siRNAs Seq2 and Seq3 after 24 hours, we observed that Seq2 and Seq3 caused the most potent downmodulation (86-95% knockdown relative to NC siRNA transfection) of all KRAS mRNAs, presumably due to the 0 mismatches of these siRNAs with all the target mRNAs. As expected, Seq2 and Seq3 also demonstrated no selectivity for oncogenic mutant KRAS mRNA over WT. In contrast, EFTX-D1 demonstrated more potent inhibition of the codon 12 and 13 mutations (G12C – 79%, G12D – 85%, G12V – 85%, and G13D – 88% knockdown relative to NC siRNA), whereas it reduced WT mRNA by only 60% and Q61H mRNA by only 49%, relative to NC siRNA transfection.

The ratios of Seq2 and Seq3 siRNA inhibition of WT mRNA to inhibition of all codon 12 and codon 13 mutants assessed are near 1 (WT:Mutant inhibition, Supplementary Fig. 2), consistent with their lack of sparing WT mRNA. Conversely, the ratios for EFTX-D1 treatments are markedly lower than 1 (G12C – 0.53, G12D – 0.37, G12V – 0.38, and G13D – 0.29), highlighting EFTX-D1’s preferential silencing of the targeted KRAS mutants relative to WT. Notably, the ratio of WT inhibition to Q61H inhibition by EFTX-D1 is near 1, which is expected as EFTX-D1 is not predicted to demonstrate preferential silencing for a codon 61 mutation.

Using the NIH/3T3 KRAS WT and G12C stably expressing cell lines, we performed a dose response experiment with EFTX-D1 and Seq2 (Fig. 3b). Whereas Seq2 demonstrated similar knockdown efficacy for both KRAS WT and G12C relative to NC siRNA across all doses tested, the knockdown efficacy of EFTX-D1 was markedly higher for G12C compared to WT at all active doses assessed.

### EFTX-D1 inhibits the KRAS mutant-induced transformed phenotype of NIH/3T3 cells

The effects of EFTX-D1 in downregulating KRAS expression and downstream MAPK activation led us to investigate the impact of EFTX-D1 treatment on KRAS-driven transformation of NIH/3T3 cells. Seq2 siRNA, which has already demonstrated robust inhibitory effects on cancer cell growth (Pecot et al. 2014) was used as a positive control in all phenotypic experiments. To observe if EFTX-D1 can reverse the KRAS oncogene-induced phenotypic loss of contact inhibition and formation of foci, we transiently transfected KRAS WT or G12C NIH/3T3 cells with EFTX-D1, Seq2 or NC siRNA. We observed that knockdown of KRAS with either EFTX-D1 or Seq2 but not NC reverted the transformed morphology of stably expressing KRAS G12C NIH/3T3 cells back towards a contact-inhibited phenotype, without affecting cells expressing WT KRAS (Fig. 4a).

**Fig. 4.**
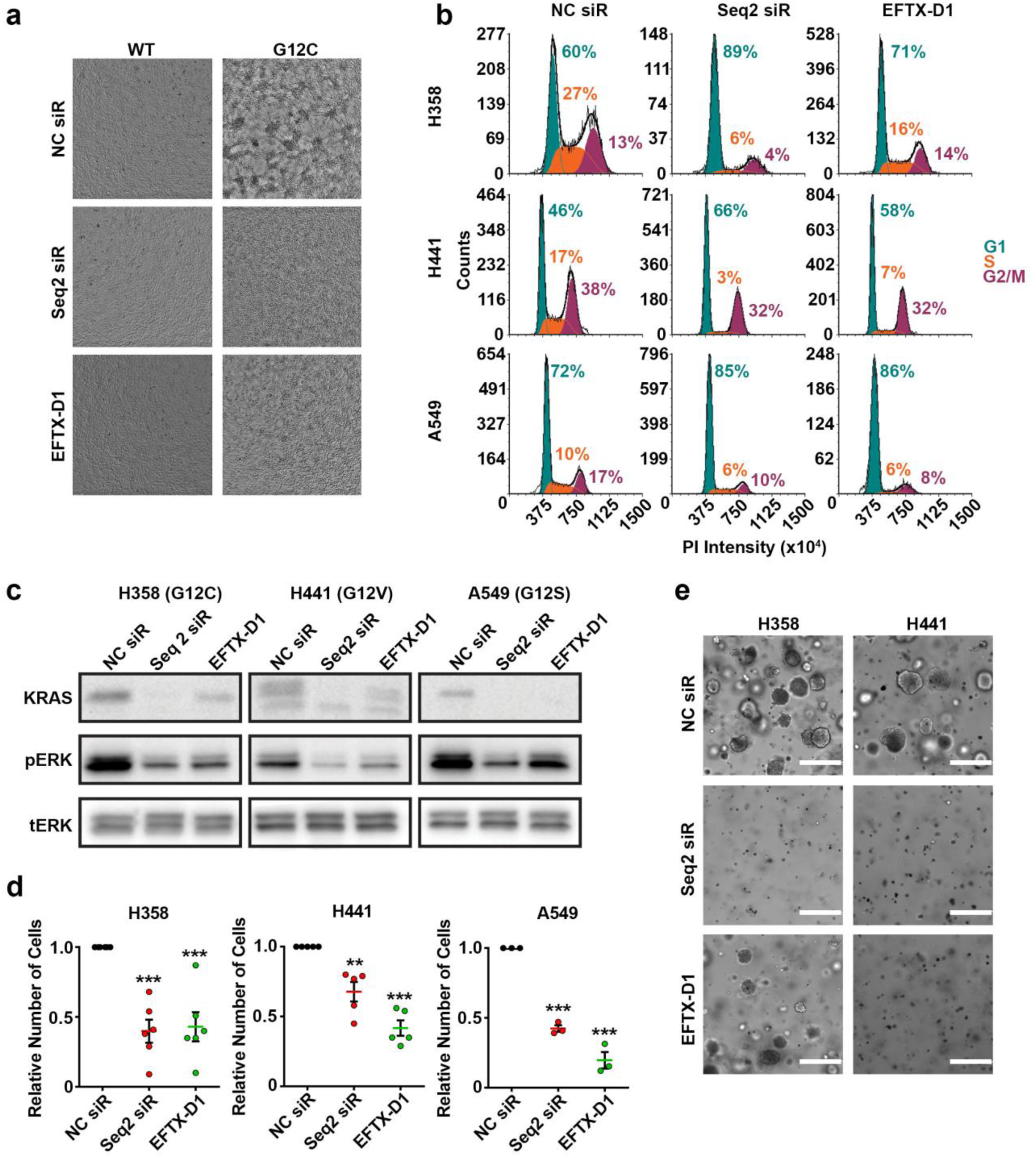
EFTX-D1 inhibits mutant KRAS-driven cancer cell growth. **a** Phase contrast images of morphological phenotypes of NIH/3T3 cells stably transduced with KRAS WT or G12C and transiently transfected with 20 nM NC, Seq2, or EFT-D1 siRNA (siR). Cells were imaged 3 days post transfection. **b** Flow cytometry cell cycle analysis of H358 (KRAS G12C), H441 (KRAS G12V), and A549 (KRAS G12V) lung cancer cells 72 hrs post transient transfection with NC, Seq2, or EFTX-D1 siR at 20 nM. **c** Western blot analysis of KRAS, pERK, and total ERK expression in H358 (KRAS G12C), H441 (KRAS G12V), and A549 (KRAS G12S) lung cancer cells transiently transfected with NC, Seq2, or EFTX-D1 siR at 20 nM. Cells were analyzed 36 hrs post transfection. **d** 2-D proliferation of H358, H441, and A549 lung cancer cells transiently transfected with NC, Seq2, or EFTX-D1 siR at 20 nM after 7 days. Samples were run in at least triplicate and cell counts are normalized to NC. **e** 3-D growth of H358 and H441 lung cancer cells embedded in Matrigel and transiently transfected with NC, Seq2, or EFTX-D1 siR at 20 nM. Cells were imaged and quantified after 7 days of growth and conditions were run in triplicate; scale bar 400 µm. Statistical significance was measured by a one-way ANOVA with a Tukey’s multiple comparisons test; p-values are indicated as **p<0.01, ***p<0.001

### EFTX-D1 inhibits KRAS-driven cancer cell proliferation

G1 cell cycle arrest is a common feature of cancer cells that have undergone a reduction in mutant KRAS. Therefore, we performed cell cycle analysis following KRAS knockdowns using EFTX-D1 or Seq2 in our panel of KRAS-dependent lung cancer cell lines harboring endogenous KRAS codon 12 mutations, including two cell lines with 3 mismatches with EFTX-D1 (H441 – G12V and A549 – G12S) (Fig. 4b). EFTX-D1 phenocopied Seq2-mediated G1 cell cycle arrest in all 3 KRAS-mutant cell lines. Treatment with EFTX-D1 and Seq2 increased the percentage of cells in G1 relative to NC siRNA treatment in all cell lines assessed: the percentage of H358 (G12C) cells in G1 increased by 29% or 11% upon Seq2 or EFTX-D1 transfection, respectively; of H441 (G12V) cells in G1 increased by 20% or 12%, respectively; and of A549 (G12S) cells in G1 increased by 13% or 14%, respectively.

As the ERK MAPK pathway is one of the major downstream effectors of KRAS signaling, we asked if EFTX-D1 treatment can also inhibit phosphorylation of ERK. Western blot analysis in H358, H441, and A549 cell lines showed a robust reduction in phosphorylated ERK upon EFTX-D1 transfection in comparison to NC siRNA transfection, similar to the effects of Seq2 (Fig. 4c).

To address if KRAS inhibition-mediated G1 arrest corresponds with reduced cellular proliferation, we characterized the effects of EFTX-D1 and Seq2 treatment on 2-D and 3-D cancer cell proliferation (Fig. 4d and e). Relative to NC siRNA treatment, EFTX-D1 and Seq2 reduced the growth on plastic of H358 cells by approximately 60% at day 7, of H441 cells by approximately 50%, and of A549 cells by approximately 25% (Fig. 4d). Finally, 20 nM of both EFTX-D1 and Seq2 also inhibited cancer cell proliferation in a Matrigel 3-D spheroid growth assay (Fig. 4e). Here, H358 and H441 cells formed spheres with an average diameter of around 100 µm while the siRNA transfection of Seq2 or EFTX-D1 inhibited the growth of spheres.

## Discussion

The KRAS oncogene has been a major target for cancer drug discovery efforts for nearly four decades. Here, we describe a novel approach to target three of the most common KRAS mutations (G12C, G12D and G13D, together comprising 57% of KRAS mutations in cancer) with one siRNA, which also preferentially spares WT KRAS. Our approach is more specific for mutant KRAS than others that impair oncogenic KRAS signaling by inhibition of key downstream kinase. The latter approach is not exclusively tailored to a cancer cell mutation, but rather to kinases that are typically ubiquitously expressed in both cancer and normal tissues. Several of these kinases, such as the kinases of the ERK MAPK pathway and the PI3K pathway, have pivotal roles in critical cellular processes, and their inhibition at higher doses or as combination therapy are reported to have side effects such as diarrhea, nausea, vomiting, reduced appetite, rash, pyrexia, fatigue and hyperglycemia {Ancuceanu, 2019 #52;Jokinen, 2015 #98}.

Thus far, the most promising direct KRAS inhibitors are targeted specifically to KRAS G12C (Canon et al. 2019; Janes et al. 2018; Ostrem et al. 2013; Hallin et al. 2020). These inhibitors covalently bind the mutated cysteine at position 12, which is not present in wild-type KRAS. KRAS G12C inhibitors are currently leading to exciting responses in clinical trials, particularly in lung cancer (Hallin et al. 2020; Canon et al. 2019), and are likely to become the first FDA-approved direct KRAS inhibitors. These successes provide proof-of-principle that direct mutant-specific inhibition of oncogenic KRAS is highly attractive and safe. However, G12C represents only 11% of all KRAS mutant cancers. An advantage of EFTX-D1 is its ability to simultaneously silence multiple KRAS G12/G13 mutations (with the most potent effects on G12C, G12D, and G13D) while partially sparing WT KRAS. This gives EFTX-D1 applicability to a much broader cohort of KRAS mutant cancers beyond only those harboring G12C, including cancers driven by KRAS G12D (33% of all KRAS mutant cancers and 19% of KRAS mutant lung cancer) and G13D (13% of all KRAS mutant cancers). In addition, targeting more than one mutation within the same tumor at the same time may help prevent the outgrowth of resistant subpopulations driven by different coexisting or *de novo* KRAS mutations {Improta, 2013 #42;Macedo, 2011 #51;Cancer Genome Atlas Research Network. Electronic address, 2017 #99}.

It is important to note that, although EFTX-D1 had the most potent binding affinity for KRAS G12C, G12D, and G13D as predicted, it still retained the capability to form a duplex with the other mutants with some level of thermodynamic stability. This suggests that there may still be sufficient interaction to enable knockdown of other mutant KRAS mRNA targets and subsequent oncogenic inhibition. This is further supported by the observation that the EFTX-D siRNAs demonstrated potent knockdown of G12V relative to WT (Fig. 1d, e), even though both G12V and WT engage in a 3-mismatch pairing with EFTX-D siRNAs (Fig. 1c). These results highlight the potential broad applicability for EFTX siRNAs to inhibit the expression of other KRAS mutants beyond G12C, G12D, G12V, and G13D, and suggest that a 3 base pair mismatch does not necessarily preclude efficacy of target knockdown. In addition to the 2 vs. 3 base pair mismatch imbalance, the position and nature of the mismatch(es) may also contribute to siRNA:target duplex thermodynamic stability and to knockdown efficacy (Du et al. 2005). For example, EFTX-D1 was able to spare WT but inhibit G12V very well, even though it has 3 mismatches with both targets. Conversely, EFTX-V1 demonstrated similar potency for inhibiting WT and G12V, although it has 3 mismatches with WT and only 2 mismatches with G12V.

Although the initial focus of the EFTX-D1 design was to reduce off-target silencing of WT KRAS in all non-tumor tissues, it is also interesting to consider effects on the silencing ratio of WT versus mutant KRAS alleles within a given cancer cell. Recent reports have suggested that wild type KRAS may harbor tumor suppressive roles in particular molecular contexts [reviewed in (Zhou, Der, and Cox 2016)]. Loss of heterozygosity of wild type KRAS has frequently been observed in the presence of mutant KRAS during tumor progression, suggesting loss of wild type KRAS may be required to enable tumor development (Li et al. 2003; Qiu et al. 2011). Similarly, restoration of wild type KRAS expression has been demonstrated to hinder tumor progression (Staffas et al. 2015). Therefore, EFTX-D1 as a therapeutic may have the added benefit of mitigating pro-tumorigenic effects of WT KRAS loss.

Over the past few years several siRNA therapeutics have received FDA approval in genetic disorders (Balwani et al. 2020; Adams et al. 2018). Recent state-of-the-art chemical modifications of siRNAs, and conjugation of siRNAs to targeting moieties, has revolutionized the field of therapeutic RNAi (Khvorova and Watts 2017). However, it remains to be seen whether nanoparticle-based or ligand-conjugated siRNA treatments will be effective in oncology. Looking forward, it is important to note that EFTX-D1 targets a region of KRAS mRNA that is fully conserved in both humans and mice; thus, future toxicity studies conducted in mice will yield information with direct relevance to human health. Taken together, our results demonstrate a novel opportunity to target oncogenic KRAS signaling and suppress cancer growth with EFTX-D1, a WT sparing, multi-mutational targeting siRNA. We believe this siRNA offers a new therapeutic tool compound in the fight against the KRAS oncogene.

## Acknowledgment

Support was provided by grants from the National Cancer Institute (NCI) to A.D.C. and/or C.J.D. from the NCI (R21CA179193, R01CA42978, R01CA175747, R01CA223775, P50CA196510, U01CA199235, P01CA203657 and R35CA232113), and from the Pancreatic Cancer Action Network/AACR (15-90-25-DER), Department of Defense (W81XWH-15-1-0611), and the Lustgarten Foundation (388222) to C.J.D. B.P. was supported by the Deutsche Forschungsgemeinschaft (DFG PA 3051/1-1). C.V.P. was supported in part by the National Institutes of Health (NIH) R01CA215075 and 1R41CA246848, a Mentored Research Scholar Grants in Applied and Clinical Research (MRSG-14-222-01-RMC) from the American Cancer Society, the Jimmy V Foundation Scholar award, the Stuart Scott V Foundation, the Lung Cancer Research Foundation, the Free to Breathe Metastasis Research Award and a North Carolina Biotechnology Translation Research Grant (NCBC TRG). S.H.A. was supported in part by an NIH 1R41CA246848 and an NCBC TRG award. P.S.P. was supported by the Volkswagen Stiftung (Project Az 92 768).

## Materials and Methods

### Cell lines and culture

NIH/3T3 mouse fibroblast cells were obtained from ATCC, and KRAS-transduced NIH/3T3 cells were maintained in Dulbecco’s Modified Eagle Medium (DMEM) supplemented with 10% Colorado calf serum, 1% penicillin-streptomycin (Pen-Strep), and 2 µg/mL puromycin. Human embryonic kidney (HEK)293T cells and A549 lung adenocarcinoma cells were obtained from ATCC. H358 bronchioalveolar carcinoma cells and H441 lung adenocarcinoma cells were obtained from NCI. HEK293T cells were maintained in DMEM; H358, H441, and A549 cells were maintained in RPMI 1640 medium. All media were supplemented with 10% FBS and 1% Pen-Strep. All cell lines were maintained in a humidified chamber with 5% CO_2_/95% air at 37°C. Cell lines were monitored for mycoplasma contamination, and all *in vitro* experiments were conducted with 60-80% confluent cultures.

NIH/3T3 cells stably expressing human KRAS WT, G12C, G12D, G12V, or G13D were generated as follows: first, retroviral particles were generated by co-transfecting 1.25 μg pBABE-puromycin retroviral vectors expressing WT KRAS or each KRAS mutant with 1.25 μg/μl PCL10A pack vector using 6.25 μg Lipofectamine 2000 (ThermoFisher Scientific) into HEK293T cells seeded in a 6 cm cell culture plate per the manufacturer’s instructions. At 24 hours post transfection, HEK293T cells were changed into fresh media. Viral supernatant was collected 24 hours later and stored on ice. Fresh media was added to the cells and again viral supernatant was collected 24 hours later. The two batches of viral supernatant were combined and filtered. NIH/3T3 cells were then seeded in a 6-well plate at a density of 100,000 cells/well and 250 μl of virus was added to the cells along with 10 μg/ml polybrene. Cells were spinoculated at 1,500 x g for 1 hour and then transduced for another 24 hours at 37°C. Virus was removed from the cells, and 24 hours later 2 μg/ml puromycin selection media was added. Cells were considered selected once all non-transduced cells in a control well were killed by the selection media.

### UV melting experiments

RNA duplexes were prepared by mixing solutions with equimolar concentrations of two strands (1.0 µM) in 1x PBS buffer, 137 mM NaCl, 2.7 mM KCl, 10 mM Na_2_HPO_4_ and 1.8 mM KH_2_PO_4_, pH 7.4. Prior to running UV melting experiments, strands were annealed by heating samples in a water bath to 70°C for 2 min, followed by slow cooling to room temperature and refrigeration overnight. All measurements were made using a Cary 100 Bio UV-Vis spectrophotometer (Agilent Technologies Inc., Santa Clara, CA), equipped with a temperature controlled multicell holder (6 x 6) and a Cary temperature controller. Absorbance versus temperature profiles were acquired by monitoring the absorbance at 260 nm (A260) between 20 and 90°C with a ramp rate of 1°C per minute. A260 values were measured at 0.2°C intervals, and melting temperatures T_m_ extracted as the maxima of the first derivatives of smoothened melting curves (filter 5) using the Cary WinUV software (Version 3.0, Agilent Technologies Inc.).

### siRNA reverse transfection and mRNA isolation

Sequences for the negative control (NC) and positive pan-KRAS control (Seq2, Seq3) siRNAs were as previously described (Pecot et al. 2014). All siRNAs were obtained from Sigma. At the time of transfection, KRAS-infected NIH/3T3 cells were plated either in 24-well plates a density of 60,000-120,000 cells per 500 μl media per well or in 12-well plates at 240,000 cells per 1 ml media per well. Cells were incubated in a 5:1 mixture of complete medium and serum-free medium, along with 20 nM siRNA (unless dose is otherwise indicated) in a 1:2 ratio with Lipofectamine RNAiMAX transfection reagent (Thermo Fisher Scientific) for 16-24 hours at 37°C in 5% CO_2_.

### Quantitative real-time PCR

For mRNA quantification, total RNA was extracted from cultured cells using the Quick RNA MiniPrep Zymo Research Kit (Genesee Scientific, #11-328) or the RNeasy Mini Kit (Qiagen, # 74104), and purified RNA was quantified using a spectrophotometer (Nanodrop). cDNA was synthesized using an iScript cDNA Synthesis Kit (Bio-Rad, #1708891) per the manufacturer’s instructions. Analysis of mRNA levels was performed on a StepOnePlus Real-Time PCR System (Applied Biosystems). Specific primers for KRAS [set #1 (forward)-TGACCTGCTGTGTCGAGAAT, (reverse)-TTGTGGACGAATATGATCCAA or set #2 (forward)-TCCAACAATAGAGGATTCCTACAG, (reverse)-CCCTCATTGCACTGTACTCCT] were used for SYBR Green-based real-time PCR, and 18S rRNA was used as a housekeeping gene. PCR was done with reverse-transcribed RNA, 1 μL each of 20 μM forward and reverse primers, and 2X PowerUp SYBR Green Master Mix (Life Technologies, #100029284) in a total volume of 25 μL. For SYBR PCR, each cycle consisted of 15 seconds of denaturation at 95°C and 1 min of annealing and extension at 60°C (40 cycles). Reactions were run in duplicate or triplicate. Relative quantitation (RQ) values were calculated using the ΔΔCt method.

### Immunoblotting

Cells were washed with ice cold PBS, lysed in RIPA buffer supplemented with phosphatase (Sigma) and protease (Roche) inhibitors, scraped, and collected in pre-chilled tubes. Lysates were then cleared by centrifugation at 15,000 x g for 10 min at 4°C, and protein concentration was determined using a Bradford assay (Bio-Rad). Standard immunoblotting procedures were followed. Membranes were blocked in Rockland blocking buffer diluted in TBS + 0.05% Tween 20 (TBST). Antibodies: Rabbit monoclonal anti-HA-Tag KRAS (Cell Signaling Technology Cat #3724), Mouse monoclonal anti-KRAS (Millipore Cat #OP24), Rabbit polyclonal anti-phospho-p44/42 MAPK (pERK1/2) (Thr202/Tyr204) (Cell Signaling Technology Cat #4370), Rabbit polyclonal anti-p44/42 MAPK (ERK1/2) (Cell Signaling Technology Cat #9102), ß-Actin (Sigma-Aldrich Cat #5441).

### Focus formation assay

3000 cells were seeded into a 96-well clear bottom plate (Corning #3904) reverse transfected as described above and imaged after 3 days with the Molecular Devices SpectraMax i3x MiniMax 300 imaging cytometer.

### 2-D Proliferation assay

3000 cells per well were seeded into Corning 96-well clear bottom plates (Corning #3904), reverse transfected as described above. After indicated days cells were stained with Calcein-AM (Invitrogen #C3100MP) and imaged with the Molecular Devices SpectraMax i3x MiniMax 300 imaging cytometer.

### 3-D proliferation assay

Cells were reversed transfected as described above and after 24 h 5000 cells were seeded into a 50 µl Matrigel (Corning) dome in a 24-well plate. After solidifying 750 µl of medium was added. Cells were cultured at 37° C in 5% CO_2_. Cells were imaged with Evos XL core.

### Cell Cycle analysis

200,000 cells were seeded into a 6 cm plate and reverse transfected with 20 nM of indicated siRNAs. After 3 days cells were washed with PBS and harvested using TrypLE. After harvesting cells, were washed once with PBS and centrifuged at 400 x g and fixed with ice cold 70% ethanol overnight. Cell pellets were resuspended in 1x PBS before adding 9 volumes of ice cold 70% ethanol dropwise and fixed overnight at 4°C. The next day, cells were washed with PBS and resuspended in cell cycle solution (1x PBS + 20 µg/ml propidium iodide and 100 µg/ml RNase) for 3 hours at 37°C. PI intensity was measured with the Beckman Coulter Cytoflex.The data was gated and analyzed with FCS Express Express 6 Flow Software by gating for cells (gate 1) and singlets (gate 2) before performing cell cycle analysis with FCS Express.

### Statistical analyses

All statistical tests were performed using GraphPad Prism 7 and 8 (GraphPad Software, Inc., San Diego, CA). All line and bar graphs represent mean values, and all error bars represent standard error of the mean.

---

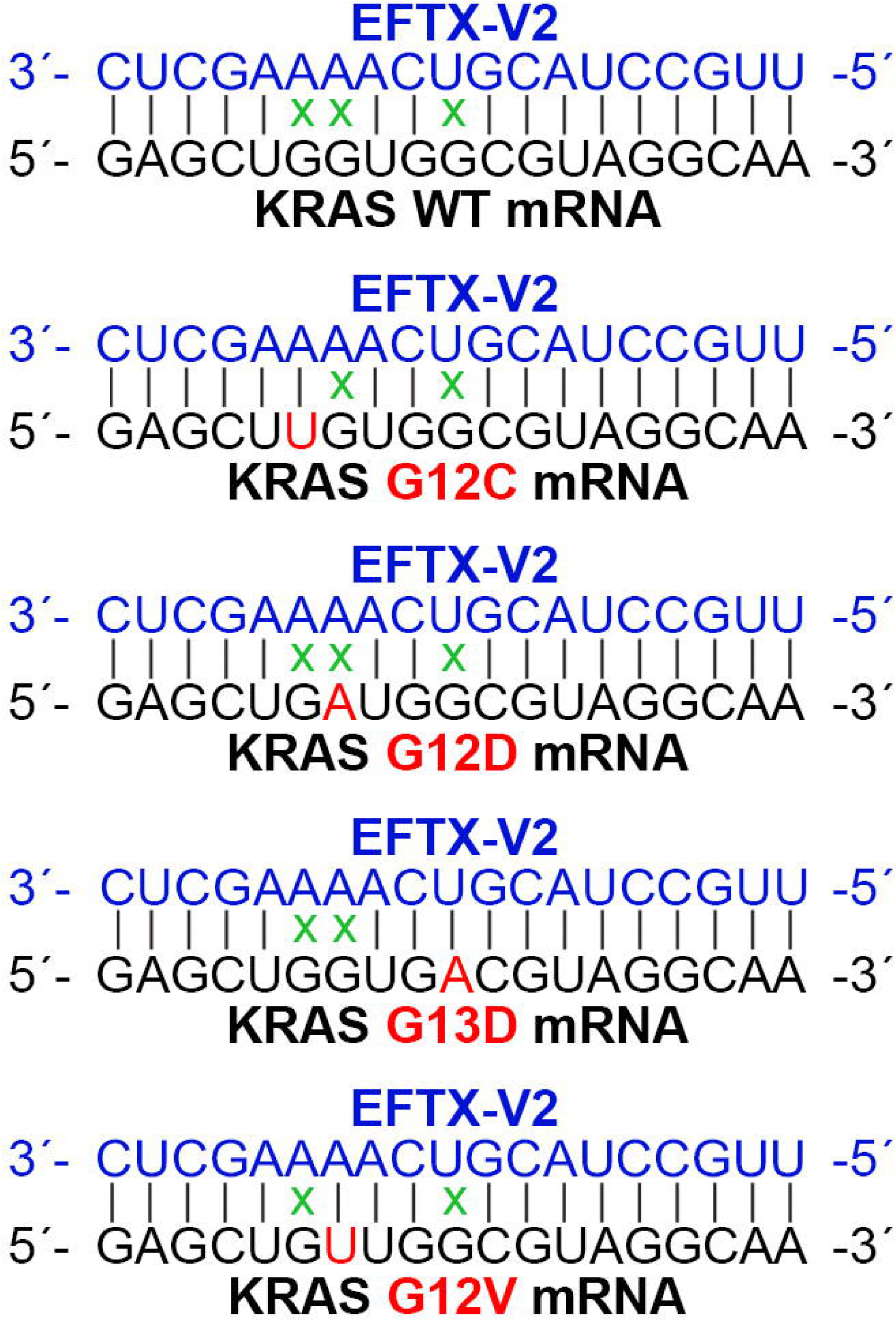

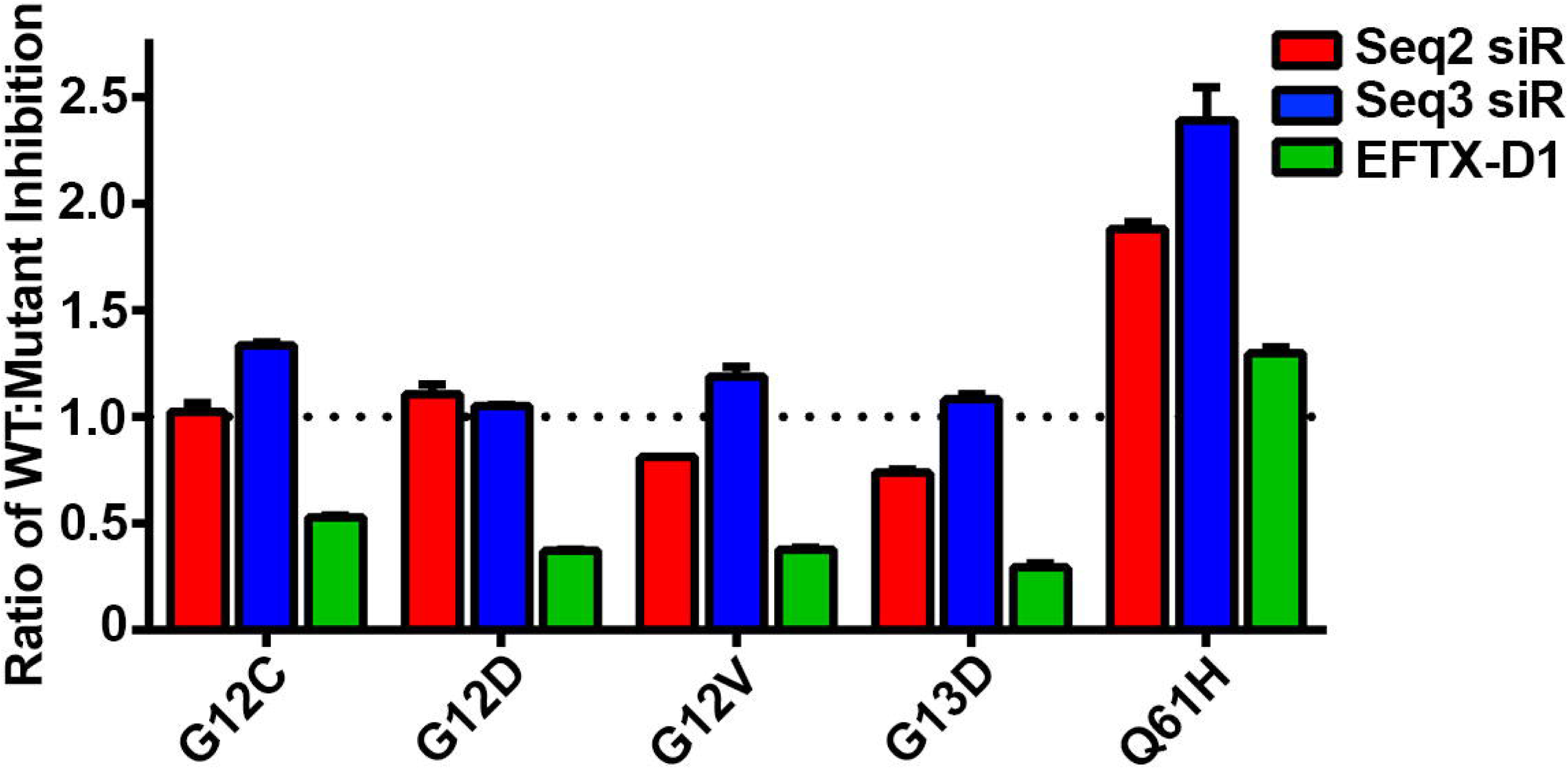

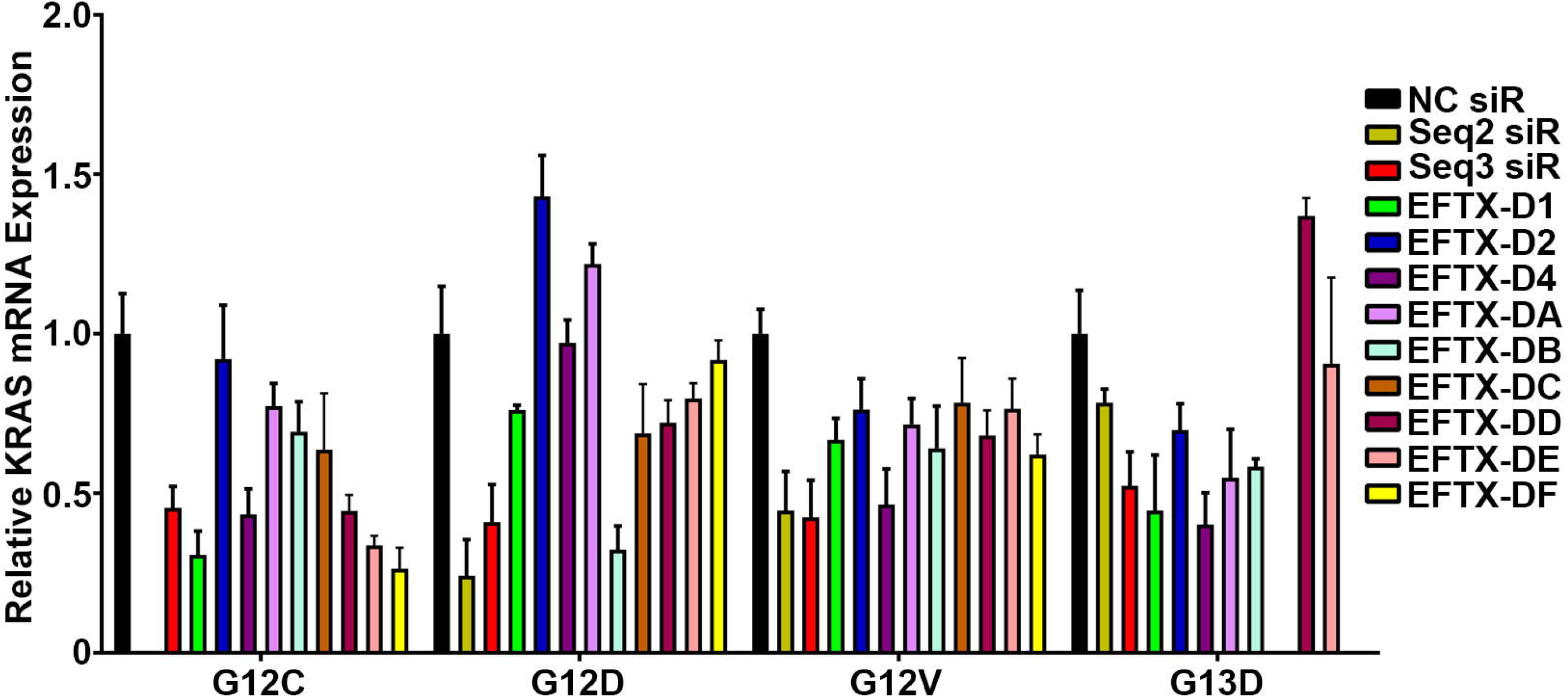

