## Supplemental Figures for "Silencing of oncogenic KRAS by mutant-selective small interfering RNA"

### Supplementary Information

Bjoern Papke, Salma H. Azam, Anne Y. Feng, Amanda E. D. Van Swearingen, Christina Gutierrez-Ford, Pradeep S. Pallan, Martin Egli, Adrienne D. Cox, Channing J. Der, Chad V. Pecot

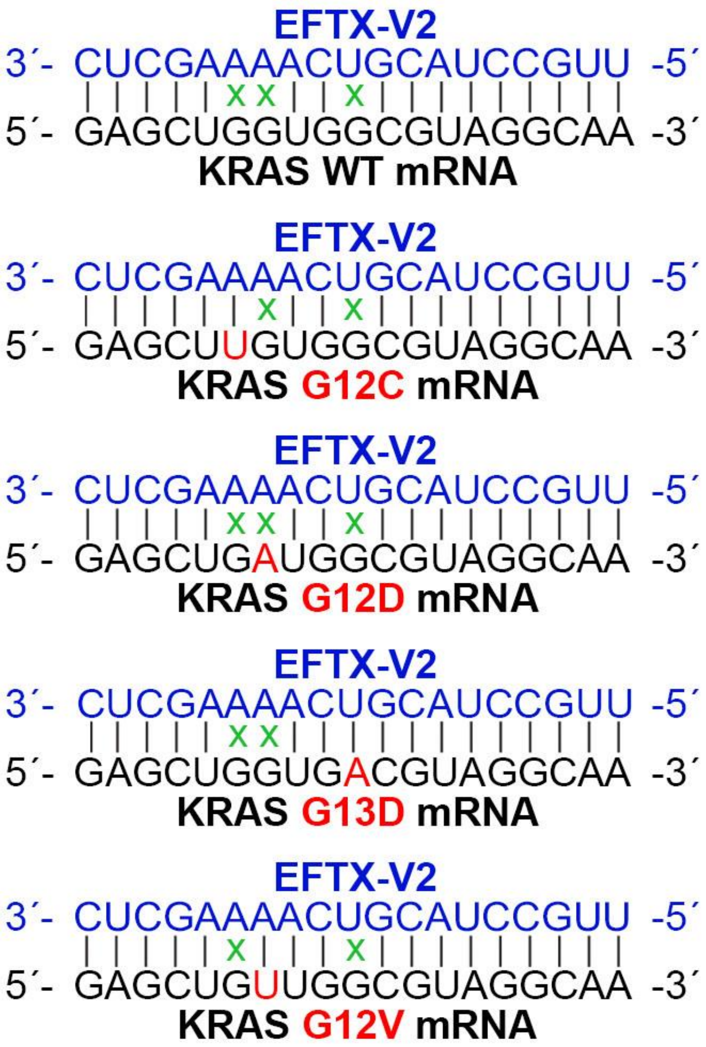


**Supplementary Fig. 1 EFTX-V siRNA targeting of KRAS transcripts.** Pairing of the EFTX-V2 antisense strand with KRAS WT, G12C, G12D, G12V, and G13D mRNA transcripts. Non-Watson-Crick base pairings (i.e., mismatches) are indicated by a green *x*.

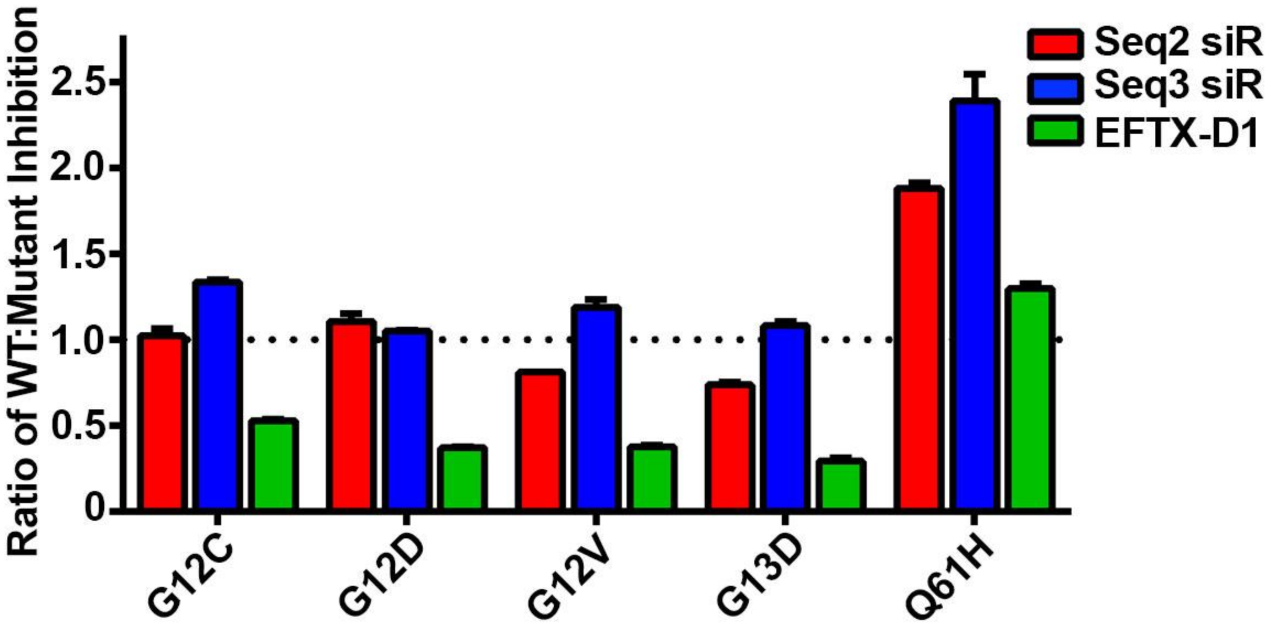


**Supplementary Fig. 2 EFTX-D1 sparing of WT.** Ratio of WT to mutant KRAS mRNA expression inhibition as determined by qPCR of NIH/3T3 cells stably transduced with KRAS WT, G12C, G12D, G12V, G13D, or Q61H and transiently transfected with negative control (NC), Seq2, Seq3, or EFTX-D1 siR at 40 nM. Cells were analyzed 24 hrs post transfection. Results were normalized to NC siR transfection (dotted black line) and RT-qPCR was run in duplicate.

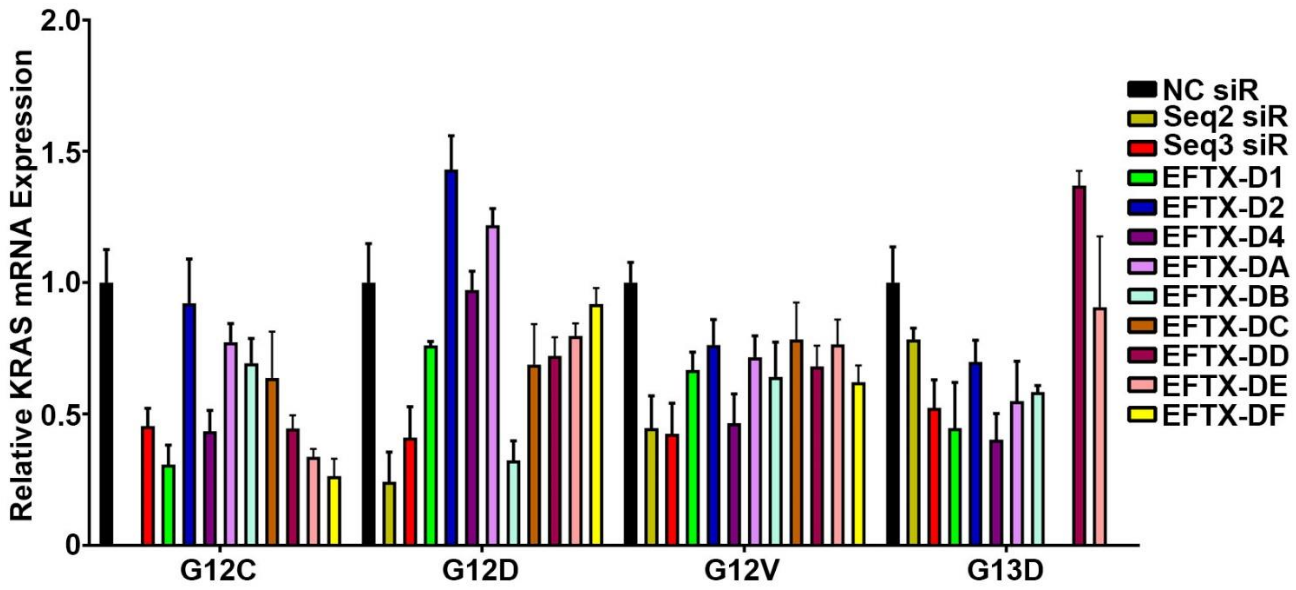


**Supplementary Fig. 3 Investigating other EFTX-D candidate siRNAs.**

qPCR of NIH/3T3 cells stably expressing KRAS G12C, G12D, G12V, or G13D and transiently transfected with negative control (NC), Seq2, Seq3, or EFTX-D siRs at a dose of 20 nM. Cells were analyzed 24 hrs post transfection and qPCR was run in triplicate. Results were normalized to NC siR transfection.
